# Estrogen receptor alpha (ERα) driven trans-regulation of mitotic checkpoint complex (MCC) components affects the clinical outcome of breast cancer

**DOI:** 10.1101/2024.08.31.610592

**Authors:** Suryendu Saha, Debanil Dhar, Stuti Roy, Ratnadip Paul, Anindya Mukhopadhyay, Arnab Gupta, Somsubhra Nath

## Abstract

Hormone receptors (HR), namely estrogen receptor (ER) and progesterone receptor (PR), are prevalent in most malignant tumors. Although previous literature provided clues for ERα in regulating mitosis and ploidy status in breast cancer (BC) cells, reports on the mitotic regulators being the targets of HR are sparse. To delve deeper into ERα’s impact on mitotic execution, our study focuses on examining its transcriptional activity on the core mitotic checkpoint complex (MCC) components, which are involved in ploidy maintenance. The expression of the core MCC components (Bub3, Mad2, and BubR1) was analyzed by quantitative-PCR and immunohistochemistry in breast tumors and adjacent normal tissues from the cancer genome atlas-breast invasive carcinoma collection (TCGA BRCA) dataset and in a prospective cohort of Eastern Indian breast cancer affected individuals. The preliminary data from these cohorts indicated an influence of ERα on the two MCC components, namely Mad2 and BubR1. Subsequently, luciferase reporter assays and chromatin immunoprecipitation were performed which revealed that ERα promotes transcriptional activation of *MAD2* and *BUB1B* through direct recruitment on these promoters, showing affects in mitotic outcome. Interestingly, the ectopic introduction of ERα, in an HR-ve breast cancer line, MDA-MB-231, significantly reduced the percent aneuploidy. Moreover, we found that overexpression of *MAD2* and *BUB1B* is associated with poorer survival in HR-positive (HR+ve) patients in both cohorts. Our findings provide insights into the specific role of ERα-mediated transcriptional regulation of mitosis and ploidy outcome. Targeting the deregulated MCC components thus offers translational potential for the therapeutic management of breast cancer.

## Introduction

Globally, breast cancer is the most commonly diagnosed malignancy [1], and the leading cause of cancer-associated death in women. This disease comprises a few clinical subtypes, based on the expression of hormone receptors (HR), namely estrogen receptor (ER) and progesterone receptor (PR), and overexpression or amplification of human epidermal growth factor receptor 2 (*HER2*) [2, 3]. Indeed, the treatment modalities in breast cancer largely vary in these receptor-defined subtypes. Around 70 to 80 percent [4] of the breast malignancies are found to be HR-positive (HR+ve) where estrogen receptor-α (ERα) and/or PR are the key drivers of carcinogenesis [5]. Such cancers are qualified for targeted treatments that block the HR activity, collectively termed endocrine therapy [6, 7]. Clinically, the HR+ve category shows lesser aggression and better prognosis than HER2-enriched (i.e. ER-ve PR-ve HER2+ve) and triple-negative breast cancer (TNBC; ER-ve PR-ve HER2-ve) subtypes, with the latter showing the poorest outcome. The prognostic consequences of breast cancer-affected individuals are deeply linked to a higher prevalence of aneuploidy, a phenotype characterized by the alteration in nuclear DNA copy numbers [8, 9]. As aneuploidy reportedly confers more aggressive traits to the tumors [10–19], it has been proposed as a prognostic marker for various malignancies [17, 20], including breast cancer [21]. It is noted that an increase in aneuploidy results in a pronounced capability to generate cellular heterogeneity; thus, the comprehensive assessment of the extent of aneuploidy and its degree might prove to be a reliable indicator of tumor aggressiveness for different cancers [15, 22]. At the molecular level, the occurrence of aneuploidy is frequently linked to improper execution of the mitotic checkpoint, also named as spindle assembly checkpoint (SAC). SAC is a feedback control mechanism that in normal cells, prevents anaphase onset until all the kinetochores complete the bipolar attachment to the mitotic spindle. This, in turn, ensures equal segregation of duplicated chromosomes into two daughter cells, preventing the rise of aneuploidy. The mechanism of SAC involves the recruitment of its effector, the mitotic-checkpoint-complex (MCC) to the kinetochore which inhibits the anaphase promoting complex/cyclosome (APC/C) activity. As an effector of the checkpoint, the MCC contains four key genes of SAC such as *MAD2L1*, *BUB1B*, *BUB3*, and *CDC20*. It prevents premature chromosomal segregation by sequestering Cdc20, a potent activator of APC/C. APC/C is a crucial E3 ubiquitin ligase whose activity is indispensable for the anaphase onset and mitotic progression of cells. Once SAC is turned off by the achievement of bipolar spindle assembly, it leads to the disintegration of MCC, which in turn, releases Cdc20, thus activating APC/C. Once activated, APC/C catalyzes the ubiquitination of two of its downstream mitotic substrates, Securin and Cyclin B1, leading to their proteasomal degradation. This event promotes anaphase entry, followed by mitotic exit [23]. The pivotal role of SAC in ensuring proper chromosomal segregation during mitosis led to the understanding that its dysfunction might be responsible for chromosomal instability (CIN) and aneuploidy. Indeed, a large number of cancer cells are found aneuploid due to defective execution of SAC [16, 24]. Additionally, studies regarding the genetic regulation of SAC components suggested a shared relationship between their altered expressions with weakened SAC, the occurrence of aneuploidy, and tumorigenesis [25–32], rather than mutations in the respective genes [33–37]. In breast cancer, SAC impairment leads to resistance against anti-microtubule drug-induced apoptosis [38]. One of the SAC components, Bub1 is reportedly overexpressed in breast cancer which is associated with cancer progression and proliferation, with a poor prognosis [39, 40]. According to another study, Mad2, one of the key MCC proteins, and its antagonist p31^comet^ were both found to be coordinately overexpressed in breast cancer. The expression of this antagonist-target pair is regulated in cells and its deregulation is correlated with cancer progression. Indeed, a lower ratio of p31^comet^ and Mad2 in breast tumors was shown to result in SAC deregulation, culminating in the generation of CIN and aneuploidy in breast cancer [41].

In ERα positive cells, Cyclin D1 (CCND1) transcription is regulated by recruitment of a coactivator complex induced by adiponectin [42, 43]. Also, breast tumors, which are typically positive for ERα, exhibit an elevated Cyclin D1 level [44, 45]. As a consequence, the *CCND1* gene is reportedly overexpressed in at least 50% of cases [46]. Indeed, the transcriptional role of ERα is directly relevant to the regulation of the cell cycle [47]. However, the information on ERα regulating the transcription of mitotic genes remains sparse. One such report showed that hormone treatment leads to upregulation of the SAC-associated gene, *ESPL1* in HR+ve breast cancer cells [48]. Furthermore, recent studies from our lab revealed that the overexpression of hormone-receptor-degrading-protein, CUEDC2, upon reducing the endogenous ERα level, brought upon a concomitant decrease of the MCC components, BubR1 and Mad2, at the transcript levels in breast cancer cell line, MCF-7. Further, this decrease was restored by a simultaneous restoration of ERα by ectopic means [49]. As the execution of SAC is largely dependent on the MCC, it seems interesting to dissect the role of the potent mitogen and the transcriptional factor, ERα, on the regulation of MCC. In this study, we aimed to decipher the molecular connection between ERα and the execution of mitosis in breast cancer. Our investigation started with assessing the expression of the MCC components in a prospective study cohort of breast cancer affected individuals from Eastern India as well as in the cancer genome atlas breast invasive carcinoma collection (TCGA BRCA) cohort. This analysis was then performed in subtype-specific breast cancer cell lines upon manipulating the level or activity of ERα. Furthermore, the study revealed a direct transcriptional role of ERα on genes coding for MCC components, Mad2 and BubR1, and the effect of this regulation on mitotic progression and ploidy outcome in breast cancer cells. Finally, we correlated this genetic regulation with the survival outcome of breast cancer study cohorts.

## Methods

### Cell culture, synchronization, transfection, and treatment

MCF-7 and MDA-MB-468 were procured from NCCS, India. T-47D and MDA-MB-231 were purchased from ATCC. Cells were cultured in DMEM (10% FBS and 1% penicillin/streptomycin), 5% CO_2,_ and 37 °C. For cell synchronization, cells were treated with thymidine (2.5 mM) for 18 hours, and released for 8 hours, with a successive second thymidine treatment for 16 hours. Post the second thymidine release (considered as ‘0’ hour) cells were treated with nocodazole (100 ng/ml) until harvest. Cells were transfected with an ERα-expressing plasmid, PCMV3-N-FLAG-ESR1 (Sino Biologicals) and a Mad2-expressing plasmid, full-length MAD2 in pDsRed1-C1 (Clontech) [50] using Lipofectamine^TM^ 2000 reagent (Invitrogen) and harvested 24-48 hours post transfection. Cells were treated with 17-β-estradiol (10nM, Sigma) for 16 hours, before harvesting. For siRNA-mediated downregulation, transfected cells were harvested after 72 hours. Mad2 siRNA was obtained from Santa Cruz Biotechnology.

### Generation of plasmid constructs

The promoter regions of *MAD2* (*MAD2L1)* and *BUB1B* were amplified from human genomic

DNA with the primers listed below.

Primers amplifying the *MAD2* promoter

F: 5’CCAGAGGGCTTAGAAGGACC3’

R: 5’TTCAAAAGTAACGACGCAGC3’

Primers amplifying the *BUB1B* promoter

F: 5’ ATCACCCCTTCCCCTCTCTC 3’

R: 5’ CCTGGGCTTTCTTCCGCAAC3’

The amplified regions were cloned into the linearized pJET1.2/blunt Cloning Vector (Thermo Fisher Scientific) according to the manufacturer’s protocol. The fragments were then subcloned into luciferase reporter vector pGL3 basic (Promega) using restriction enzyme BglII (New England Biolabs). The deletion constructs of *MAD2* as well as *BUB1B* promoters were generated by omitting one of the estrogen response element (ERE) using Quikchange II site-directed mutagenesis kit (Agilent).

### TCGA data retrieval and gene expression analysis

The gene expression data of the TCGA BRCA dataset were extracted from the UALCAN database (https://ualcan.path.uab.edu/). RNA-seq data were retrieved in the form of transcripts per million (TPM) values for three MCC component genes, namely *BUB3*, *MAD2*, *BUB1B*. A total of 1211 samples, of which 1097 were from the primary breast tumor origin and 114 from adjacent normal tissues, was included in the present study. Adjusted p value < 0.05 was used to identify the differentially expressed genes.

### Cancer Cell Line Encyclopedia (CCLE) data retrieval and analysis

The gene expression data of two of the MCC genes (*MAD2* and *BUB1B*) were extracted from the 22Q2 public data repository from the DepMap portal at Broad Institute (https://depmap.org/portal/). The mRNA expression data [Log2 (TPM+1) values] of the HR+ve breast cancer cell lines were acquired and Pearson’s correlation analysis was performed.

### Western Blotting (WB)

Cells were lysed using NP-40 lysis buffer (Amresco) and protein concentration was estimated by Bradford’s assay (Biorad Laboratories Inc.). Equal amounts of protein were resolved using SDS-PAGE, transferred to PVDF membrane (Amersham), and incubated overnight in primary antibodies against Bub3 (ab133699), Mad2 (ab70385), β-actin (ab8227) (Abcam), ERα (8644S), Cyclin B1 (4831S) (Cell Signaling Technology), and RFP (NBP1-96752) (Novus Biologicals). Membranes were then incubated in HRP-conjugated secondary antibody (Sigma). The immuno-complexes were visualized using chemiluminescence (Super signal West Pico, Thermo Fisher Scientific).

### Tissue specimens

Primary malignant breast tumor and adjacent normal tissues were collected with signed consent from the participants at Saroj Gupta Cancer Centre and Research Institute (SGCCRI), Kolkata. The study was approved by Institutional Ethics Committee of SGCCRI, under regulation of the Govt. of India (Registration No. ECR/250/Inst/WB/2013/RR-20, on 27^th^ July, 2020). The clinico-pathological data and follow-up information of the recruited cohort were recorded.

### Quantitative Real Time (qRT) PCR

Total RNA was extracted from cell lines and breast tissue samples using the TRIzol (Invitrogen, Thermo Fisher Scientific Inc.) reagent. Thousand ng of total RNA was used to generate cDNA using M-MuLV reverse transcriptase (New England Biolabs). Real-time PCR was performed using PowerUp SYBR™ Green master mix (Thermo Fisher Scientific Inc.) in the 7500 Real-Time PCR instrument (Thermo Fisher Scientific Inc.). Data were normalized using the expression of 18S rRNA. Log transformed values of Delta C_t_ and fold changes (wherever applicable) were plotted and statistically analyzed using unpaired t-test or Mann-Whitney U-test. All the experiments were performed in triplicate. Primer sequences used for qRT-PCR are: ESR1-5’GGCTACATCATCTCGGTTCC3’ (F), 5’TCAGGGTGCTGGACAGAAA3’ (R); BUBR1-5’AGCCAGAACAGAGGACTCCA3’ (F), 5’TGAAGCTGTATTGCCACGAG3’ (R); MAD2-5’GTGGTGAGGTCCTGGAAAGA3’ (F), 5’CCGACTCTTCCCATTTTTCA3’ (R) BUB3-5’CTACGATCCAACGCATGCCTG3’(F), 5’CAGCCGGTCTCCAGACACTGAG3’ (R) 18S rRNA-5’TGACTCTAGATAACCTCG3’ (F), 5’GACTCATTCCAATTACAG3’ (R)

### Immunohistochemistry (IHC)

Formalin-fixed paraffin-embedded breast tissues (tumor and adjacent normal) were cut into 5μm sections. The sections were then baked at 65°C for 20 minutes, followed by de-paraffinisation with two subsequent changes of xylene and rehydration in decreasing grades of ethanol. Heat mediated antigen retrieval was performed using citrate buffer (10mM sodium citrate, 0.5% Tween 20, pH-6.0) and endogenous peroxidase was quenched using 0.3% H_2_O_2_ (Merck). The sections were then incubated in primary antibodies against Bub3 (ab133699) (Abcam), Mad2 (sc-47747) (Santa-Cruz) and BubR1 (ab70544) (Abcam) at a dilution of 1:100 for overnight, followed by incubation in HRP-conjugated secondary antibody (Sigma) for 2 hours. The complexes formed by the immuno reaction were visualized by addition of 3,3’-Diaminobenzidine (Sigma), while the nuclei were stained with Mayer’s hematoxylin (Merck). Following this, the sections were dehydrated using a series of ethanol solutions and mounted using Dibutylphthalate Polystyrene Xylene (DPX) (Merck). Images were captured at a magnification of 40X using Lawrence and Mayo’s LM-52-1704 light microscope, with image acquisition performed using ScopeImage 9.0 software.

### Fluorescence *in situ* hybridisation (FISH)

Cultured cells were harvested in 1X phosphate-buffered saline (PBS). Cells were then incubated in 1:1 mixture of 0.4% KCl and 0.8% sodium citrate for 20 min at 37°C, followed by fixation in 3:1 mixture of methanol and acetic acid for 20 min on ice. Fixed cells were overlaid on charged slides and baked at 56°C overnight. On the following day, cells were dehydrated in a graded manner and subsequently incubated overnight at 42°C with denatured probes against centromere of chromosome 1. The next day, slides were washed with successive low- and high-salt buffers and nuclei were counterstained using 4′,6-diamidino-2-phenylindole (DAPI). Fluorescent signals were then visualized and enumerated at 40X magnification under fluorescence microscope (Zeiss Axio Scope.A1). The percent aneuploidy was calculated using the following formula.

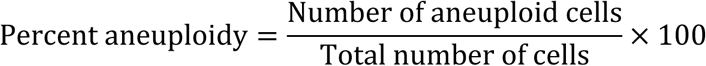

### Luciferase Reporter Assay

After transfection and/or treatment, cells were washed with 1X PBS and subsequently lysed with 5X passive lysis buffer supplied with the luciferase assay kit (Promega). Following a brief vortex, the complete cell lysates were subjected to centrifugation at 4 °C and 13,000 rpm for a duration of 2 minutes. Subsequently, 15-30 µl of the resulting supernatant was combined with 30-60 µl of luciferase assay substrate. Luminescence was measured as relative luciferase unit in a GLOMAX luminometer (Promega). Total protein concentration in each lysate was measured by protein assay reagent (Bio-Rad) and then used to normalize the luciferase activity of each lysate. Each assay was performed in duplicate and repeated three times. Fold activation values were calculated as mean of three separate experiments.

### Chromatin Immunoprecipitation (ChIP) Assay

After transfection, cells were first cross-linked with 37% formaldehyde (Sigma). Then the process was stopped by adding 1.25M glycine (Thermo Fisher Scientific Inc.). Next, cells were washed with 1X PBS and lysed in SDS lysis buffer (50mM Tris-HCL, pH 8.0, 10mM EDTA, 1% SDS) and subjected to sonication. Sonicated genomic DNA was diluted with ChIP dilution buffer (0.01% SDS, 1.1% Triton X 100, 1.2mM EDTA, 16.7mM Tris-HCL, pH 8.1, 167mM NaCl). Immunoprecipitation was executed using primary antibody against ERα (8644S, Cell Signaling Technology), or normal IgG control (Sigma) at 4°C overnight. The antibody-protein complex was pulled down using Protein G sepharose beads (Thermo Fisher Scientific). The beads were washed with low salt immune complex buffer (0.1% SDS, 1% Triton X 100, 2mM EDTA, 20mM Tris-HCL, pH 8.1, 150mM NaCl), followed by high salt immune complex buffer (0.1% SDS, 1% Triton X 100, 2mM EDTA, 20mM Tris-HCL, pH 8.1, 500mM NaCl), LiCl immune complex buffer (0.25M LiCl, 1% IGEPAL CA630, 1% deoxycholic acid, 1mM EDTA, 10mM Tris, pH 8.1), and finally, TE buffer (10mM Tris-HCL, pH 8.0, 1mM EDTA). Chelex-100 was added to the beads and incubated at 100°C. Proteinase K and RNaseA was used for degrading associated protein, antibodies and RNA. Following this, DNA was extracted by phenol-chloroform method. qRT-PCR amplification of precipitated chromatin was done using sets of primers encompassing *MAD2* and *BUB1B* promoter as mentioned below.

Primer sequences encompassing *MAD2* promoter:

(F): 5’ TGTGACTGAGCAAAACCG 3’

(R): 5’ AAACTTTCATGTCTGCCGGA 3’

Primer sequences encompassing *BUB1B* promoter:

(F): 5’ GGACTGAGCAGAAGAGAAAGC 3’

(R): 5’ GCGTGACTCTTGTAGGGTGA 3’

### Assessment of clinicopathological features in TCGA BRCA and Eastern Indian cohort

TCGA BRCA cohort: The gene expression data of *MAD2* and *BUB1B* from HR+ve background initially composed of 911 records and its associated clinicopathological data were retrieved from the TCGA BRCA dataset in UCSC Xena (https://xenabrowser.net/). The downloaded dataset had the following variables, transcripts per million (TPM) values of *MAD2* and *BUB1B*, tumor stage according to AJCC, tumor node status, and, tumor histology based on ICD-O-3 classification, overall survival time, and recurrence free survival time. Gene expression levels of MCC genes were further segregated into high and low groups based on their median value. The categorical data were represented as percentages. After filtering out the files with missing variables, the final number of remaining case records was 581. The association between *MAD2* or *BUB1B* levels and clinicopathological variables was examined by Chi-square test for univariate analysis. Gene expression data of *MKI67* (gene name for Ki-67) was also extracted from the TCGA BRCA dataset, using UCSC Xena (https://xenabrowser.net/), taking only HR+ve cases for the downstream analysis. P-values less than 0.05 at a two-sided level were considered significant. Data management was performed using Microsoft Excel, and statistical analysis was conducted using GraphPad PRISM V8 software. Moreover, the variables were incorporated into a Cox regression model, employing IBM SPSS software (IBM Corp. Released 2013. IBM SPSS Statistics for Windows, Version 22.0. Armonk, NY: IBM Corp.), to delve into their potential predictive significance for overall survival (OS) and recurrence free survival (RFS). A multivariate Cox regression analysis was conducted using the backward conditional method, encompassing the subsequent variables: tumor stage, categorized as III/IV vs I/II (with I/II as the reference category); Nodal Involvement, categorized as N2+N3 vs N0+N1 (with N0+N1 as the reference category); Ki67 gene expression, categorized as ‘High’ vs ‘Low’ (with low as the reference category); histological type, categorized as infiltrating ductal carcinoma (IDC) vs others (with others as the reference category); *BUB1B* expression, categorized as ‘High’ vs ‘Low’ (with Low as the reference category); *MAD2* expression, categorized as ‘High’ vs ‘Low’ (with Low as the reference category). A p-value below 0.05 was considered as statistically significant.

Eastern Indian cohort: A total of 46 HR+ve breast cancer-affected individuals from Eastern India was included in this study. The clinicopathological characteristics of the cohort were recorded. The gene expression levels of *MAD2* and *BUB1B* were assessed using qPCR. The generated dataset was divided into two groups (‘High’ and ‘Low’), based on the median expression of *MAD2* or *BUB1B*. Univariate parameters, including tumor stage and grade, lymph node involvement, proliferation index (Ki-67), and histologic types were considered for statistical analyses. Data management was performed using Microsoft Excel, and statistical analysis was conducted using GraphPad PRISM V8 software. Association between *MAD2* or *BUB1B* levels and clinico-pathological variables were examined by Chi-square test for univariate analysis. P-values less than 0.05 at a two-sided level was considered significant. However, some of the data for different clinical parameters were not available. Therefore, cases with missing data were excluded from the dataset analyzed for those specific parameters.

### Survival analysis by KM-Plotter

Patients were divided into higher and lower expressing groups, according to the auto-computed best cutoff of mRNA expression, and the number of risk cases was verified by the Kaplan-Meier survival curve. The hazard ratio (HR) with 95% confidence intervals and log-rank *p*-value were computed. The Kaplan-Meier plotter database (http://kmplot.com/analysis/) site was used to validate the prognosis (OS and RFS) of high and low-expressing groups of *MAD2* and *BUB1B* in HR+ve breast cancer by using the ‘203362_s_at’ and ‘203755_at’ datasets, respectively. A univariate Cox *P* < 0.05 was defined as statistically significant.

### Colony formation assay

Cells, at a number of 10^3^ per 35 mm^2^ of the surface area of the culture dish, were seeded in triplicate for both the control and *MAD2* overexpression sets and incubated to allow the formation of colonies. Following this, the dishes were washed with 1X ice-cold PBS, and fixed in a solution of acetic acid and methanol (1:7). Colonies were stained in 0.1% crystal violet solution and subsequently counted.

### Statistical analyses

All the relevant experiments were carried out in triplicate and the data were analyzed using an unpaired t-test using GraphPad Prism version 8.0. The Pearson correlation analysis was performed to examine the relationship among the expression of MCC genes. The association between *MAD2* or *BUB1B* levels and clinicopathological variables was examined by Chi-square test for univariate analysis. Survival parameters were calculated by Kaplan–Meier analysis using IBM SPSS software (IBM Corp. Released 2013. IBM SPSS Statistics for Windows, Version 22.0. Armonk, NY: IBM Corp.).

## Results

### The expression of MCC components, Mad2 and BubR1, is regulated in an ERα-dependent manner in breast cancer

Initially, we analyzed the TCGA BRCA dataset and observed an upregulated transcript level of MCC components, namely *BUB3*, *MAD2* and *BUB1B* (coding for BubR1 protein) (Fig. 1A). We extended this observation in a prospective study cohort comprising Eastern Indian cases of breast cancer affected individuals. Here too, we found a similar observation for *MAD2* and *BUB1B* (Fig. 1B). Following this, we grouped the cases of this prospective cohort into ‘HR+ve’ and ‘HR-ve’ subtypes. The real-time PCR data showed a significantly upregulated *MAD2* transcript level in primary malignant breast tissues of the ‘HR+ve’ subtype, as compared to the ‘HR-ve’ tumors (Fig. 1C). Immunohistochemistry further corroborated this set of observations for Mad2 expression (Fig. 1D). Although, both the transcript levels of *BUB3* and *BUB1B* did not show any significant differences between these two subtypes (Fig. 1C), immuno-histochemistry analyses showed higher expression of BubR1 in ‘HR+ve’ tumors, in comparison to the ‘HR-ve’ subtype (Fig. 1D). We extended our observations using cultured breast cancer lines. We used two cell lines from HR+ve background (MCF-7 and T-47D) and two from HR-ve background (MDA-MB-231 and MDA-MB-468). The Western blot analyses revealed higher levels of Bub3 and Mad2 expression in both MCF-7 and T-47D, as compared to the two HR-ve breast cancer lines (Fig. 1E).

**Figure 1:**
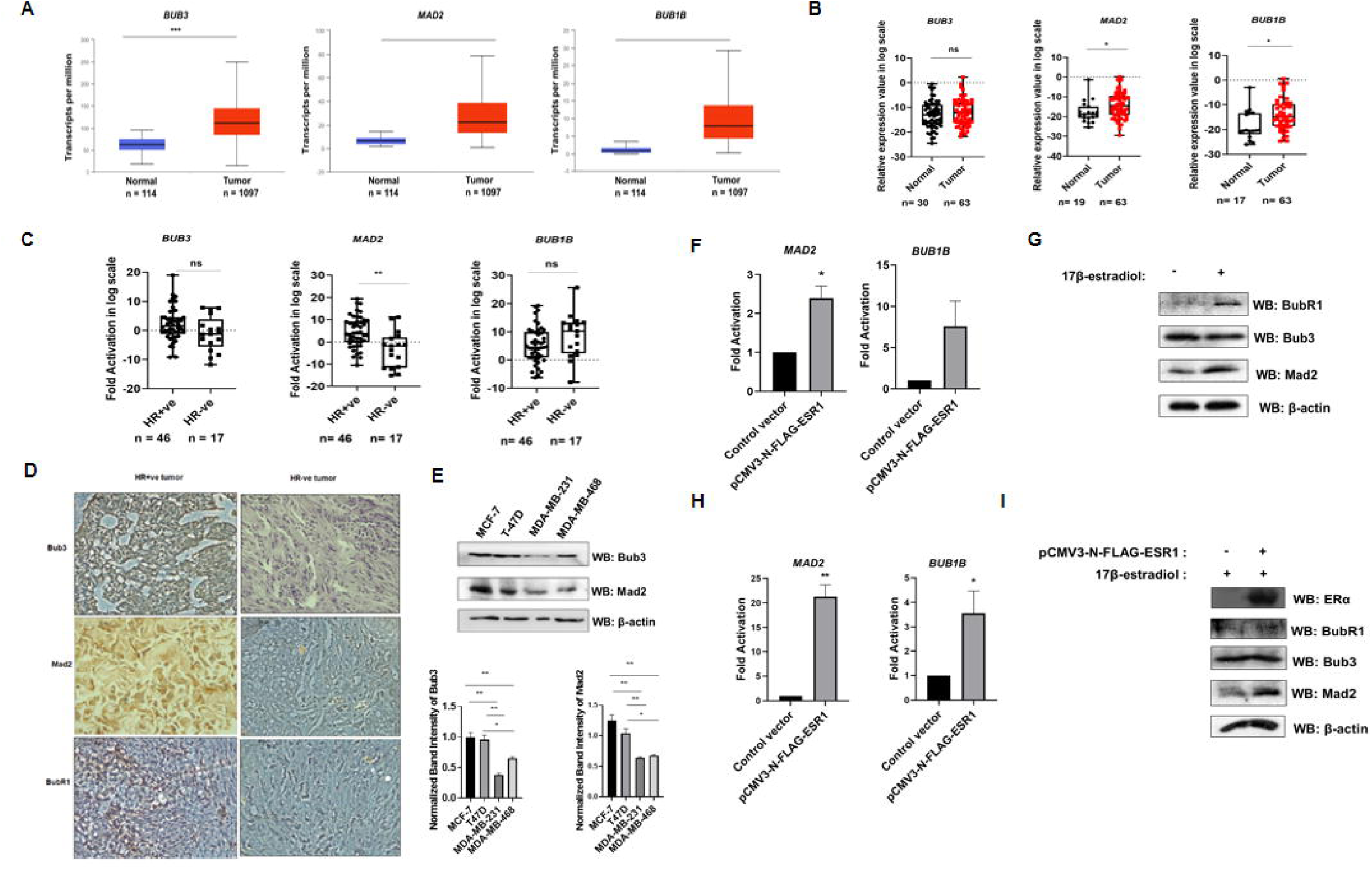
ERα upregulates the expression of MCC proteins, Mad2 and BubR1 in breast cancer. **(A)** Box plots, derived from UALCAN, representing mRNA levels of *BUB3, MAD2,* and *BUB1B* across the breast tumors (n=1097) and adjacent normal tissues (n=114) in TCGA BRCA (the cancer genome atlas-breast invasive carcinoma) dataset. ***p < 0.0005. The Y axis represents transcripts per million (TPM) for gene expression. **(B)** The expression of MCC genes in primary breast tumor tissues (n=63) and adjacent normal tissues from the prospective study cohort of Eastern Indian individuals. qRT-PCR was performed by using primers specific for *BUB3, MAD2, BUB1B* and 18S rRNA as endogenous control. The x-axis of the box plot shows the number of breast tumors and normal samples, and the y-axis shows the relative expression in the log scale. Data are represented as Box-Whisker plots. *p < 0.05; ns, not significant (Mann-Whitney U-Test). **(C)** Dot-plot representing the expression of *BUB3, MAD2,* and *BUB1B* between HR+ve and HR-ve cases of the prospective study cohort. qRT-PCR was performed as mentioned in (B). The x-axis shows the number of breast tumor cases of HR+ve and HR-ve subtypes, and the y-axis shows the fold activation in the log scale. The data were analyzed by the Mann-Whitney U-Test. **p < 0.005; ns, not significant. **(D)** Representative images of immunohistochemical analysis of HR+ve and HR-ve of primary breast malignancies using antibodies against Bub3, Mad2, and, BubR1. **(E)** Western blot showing the endogenous levels of Bub3 and Mad2 in HR+ve (MCF-7 and T-47D) and HR-ve (MDA-MB-231 and MDA-MB-468) breast cancer cell lines. Total cell lysates were prepared and a Western blot was performed using antibodies specific for Bub3, Mad2, and, β-actin (left panel). The histograms show the normalized band intensity of Bub3 and Mad2 (right panel). *p < 0.05; **p < 0.005; (two-tailed unpaired Student’s t-test). **(F)** Histograms depicting the ectopic ERα-mediated fold activation of *MAD2,* and *BUB1B* transcripts in MCF-7 cells. qRT-PCR was performed by using primers specific for *MAD2, BUB1B,* and 18S rRNA as endogenous control. The y-axis of the histograms shows the fold activation. *p < 0.05; ns, not significant (two-tailed unpaired Student’s t-test) **(G)** Western blot representing the differential expression of MCC proteins (BubR1, Bub3, and, Mad2) upon E2 (17β-estradiol) stimulus. MCF-7 cells were treated with 17β-estradiol for 16 hours. Total cell lysates were processed for Western blot by using antibodies specific for BubR1, Bub3, Mad2, and, β-actin. **(H-I)** Ectopic ERα upregulates the MCC components in (H) transcript as well as (I) in protein levels in MDA-MB-231 cells. Cells were transfected with 1.5 µg of ectopic ERα (pCMV3-N-FLAG-ESR1). qRT-PCR and western blot were performed as mentioned in (F) and (G), respectively. The y-axis of the histogram (H) shows the fold activation. **p < 0.005; *p < 0.05; (two-tailed unpaired Student’s t-test). All the experiments were performed in triplicates and error bars were represented as means ± s.d.

To investigate the potential involvement of HR on the expression of MCC components, the transient transfection of ERα in the HR+ve breast cancer line, MCF-7 showed a sharp increase in the transcript levels of both *MAD2* and *BUB1B* (Fig. 1F). These cells were also treated with a derivative of estrogen, 17-β-estradiol (E2) which stimulates the activity of ERα. Through WB analyses, we could find an upregulated expression of Mad2 and BubR1 upon this treatment (Fig. 1G). Furthermore, the ectopic introduction of ERα and concomitant E2 treatment in the cells from an HR-ve breast cancer cell line, MDA-MB-231, resulted in an increase of both *MAD2* and *BUB1B* at the transcript level (Fig. 1H). Although in the case of Mad2, there was an increased expression at the protein level, we could find no such visible change in the case of BubR1 (Fig. 1I). Similarly, for the MCC component, Bub3, no such observations were noticed in any of the above experiments (see Fig. 1G and 1I).

### ER**α** transcriptionally regulates Mad2 and BubR1 expression in breast cancer cells

The upregulation of the transcripts of the MCC components, Mad2 and BubR1, under the influence of ERα, led us to search for the putative estrogen response elements (EREs) in the promoter region of these genes by *in silico* analysis (Fig. 2A). The *BUB1B* and *MAD2* promoters, both, showed ERE sites; however, no such site could be found in the promoter region of *BUB3*. To address whether ERα is directly involved in regulating the expression of these MCC genes, we cloned the corresponding promoter regions of these two genes [an 1128-bp region of the *MAD2* (−1098 to +30 nt, named as ‘pSS-MAD2’) and a 1546-bp region of the *BUB1B* (−1454 to +92 nt, named as ‘pSS-BUB1B’)] into a reporter vector, pGL3 basic, and proceeded for luciferase assay. The estrogen treatment resulted in a significant increase in the luciferase activity from both the pSS-BUB1B and pSS-MAD2 constructs when transfected in the HR+ve breast cancer line, MCF-7 (Fig. 2B). Another cell line from the HR+ve background, T-47D, showed the similar observation (Supplementary file 1). For further validation, we extended this finding upon co-transfection of pSS-BUB1B or pSS-MAD2 with ERα expression vector, pCMV-N-FLAG-ESR1, in HR-ve MDA-MB-231 cells. The results showed a significant increase of luciferase activity upon the introduction of ERα, which was further augmented with a simultaneous E2 treatment (Fig. 2C).

**Figure 2:**
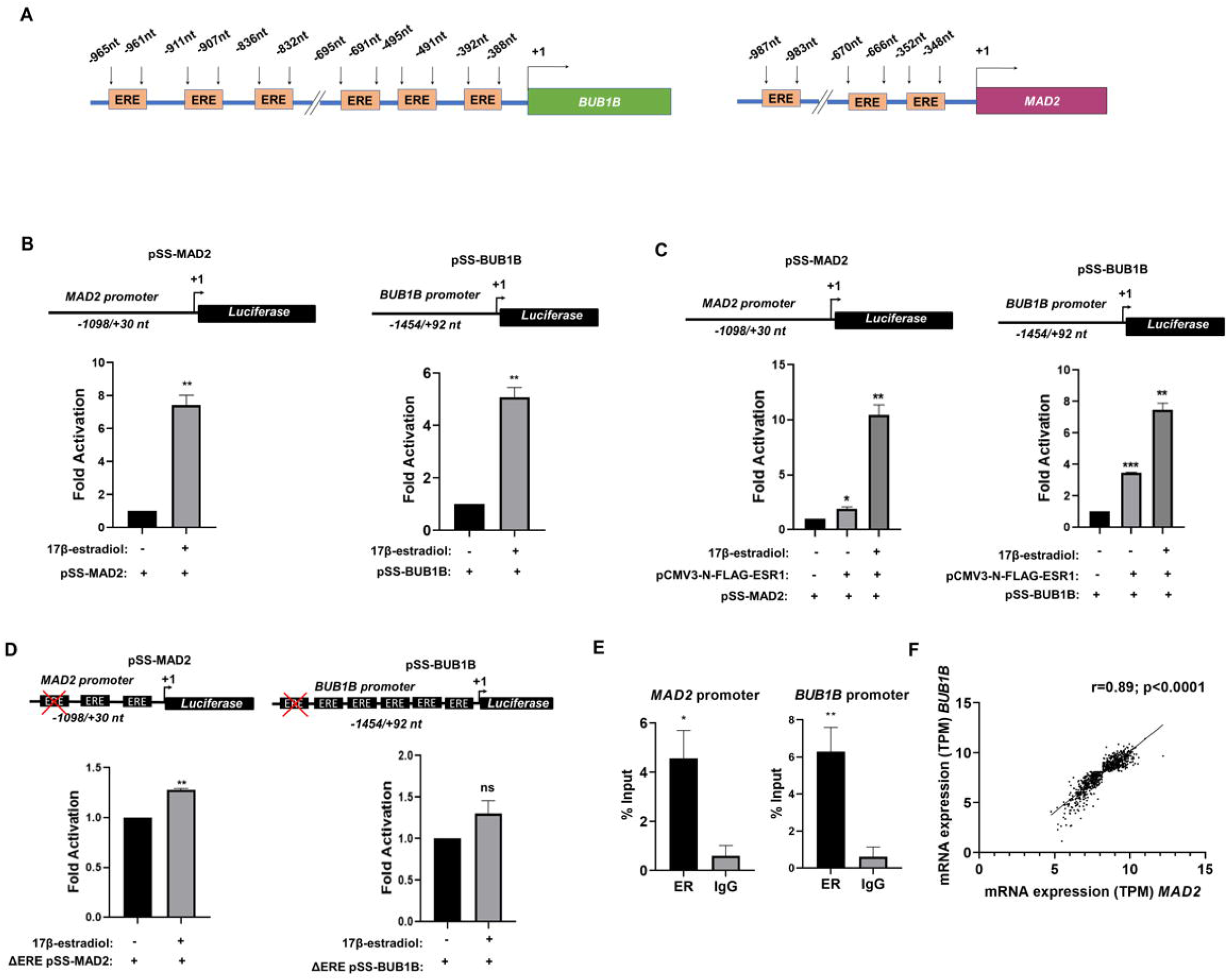
*MAD2* and *BUB1B* are transcriptionally activated by ERα. **(A)** Schematic representations showing the estrogen response elements (EREs) in the promoter regions of *MAD2* and *BUB1B*. **(B)** E2 (17β-estradiol) treatment up-regulates *MAD2* and *BUB1B* promoter activity in MCF-7 cells. The top panel shows the map of *MAD2* and *BUB1B* promoter regions used for luciferase reporter assay. pSS-MAD2 construct contains an 1128-bp region (−1098 to +30 nt) of the *MAD2* gene and the pSS-BUB1B construct contains a 1546-bp region (−1454 to +92 nt) of the *BUB1B* gene, cloned into pGL3 basic vector. MCF-7 cells were transiently transfected with pSS-MAD2 (250 ng) and pSS-BUB1B (500 ng), followed by E2 treatment, and proceeded for luciferase reporter assay. **p < 0.005; (two-tailed unpaired Student’s t-test) **(C)** Ectopic ERα up-regulates *MAD2* and *BUB1B* promoter activity in MDA-MB-231 cells. MDA-MB-231 cells were co-transfected with pSS-MAD2 (250 ng) and pSS-BUB1B (500 ng) along with 1.5 µg of ectopic ERα (pCMV3-N-FLAG-ESR1), followed by E2 treatment and proceeded for luciferase reporter assay and plotted as discussed in (B). *p < 0.05; **p < 0.005; ***p < 0.0005 (two-tailed unpaired Student’s t-test). **(D)** Deletion of ERE results in a significant decrease in luciferase activity. Map of the two deletion constructs, ΔERE pSS-MAD2, and ΔERE pSS-BUB1B were shown in the top panel. MCF-7 cells were transfected with either ΔERE pSS-MAD2 (250 ng) or ΔERE pSS-BUB1B (500 ng), followed by E2 treatment. Data were plotted as discussed in (B). **p < 0.005; ns, not significant (two-tailed unpaired Student’s t-test). **(E)** ERα physically recruits to *MAD2* and *BUB1B* promoters. The total cell extract of MCF-7 was prepared and Chromatin immunoprecipitation was done using antibodies specific for ERα and normal IgG as control. Precipitated chromatin was estimated by qRT-PCR. The results are expressed as a percent of input. *p < 0.05; **p < 0.005; (two-tailed unpaired Student’s t-test). **(F)** Representative scatter plot depicting the Pearson’s correlation analysis between the gene expression values (TPM) of *MAD2* and *BUB1B*, extracted from TCGA BRCA cohort using UCSC Xena database. The Pearson correlation coefficient is reported as the ‘r’ value. **\*\***\*p < 0.0005; (two-tailed unpaired Student’s t-test). All the experiments (B-E) were performed in triplicates and error bars were represented as means ± s.d.

However, upon obliterating the most distant ERE site from either *MAD2* or *BUB1B* promoter regions, the estrogen treatment could not reproduce the similar fold change (Fig. 2D), as compared to the wild type promoter constructs shown in Figure 2B. Distant ERE sites have been reported to act as enhancers for estrogen-responsive genes [51]. Our findings suggest that the distant EREs in both *MAD2* or *BUB1B* promoter regions could potentially act as enhancers. These data provide a supportive clue that ERα might get physically recruited on both the *MAD2* and *BUB1B* promoters and regulate their transcription. To check that functionality, we performed chromatin immunoprecipitation (ChIP) analysis followed by qPCR using the primers encompassing the specified promoter regions of *MAD2* and *BUB1B* in the MCF-7 cells. The data revealed the physical recruitment of endogenous ERα on the promoters of both the genes (Fig. 2E). Additionally, we performed Pearson’s correlation analysis between the expressions of these two genes in HR+ve cases from the TCGA BRCA cohort. This analysis revealed a positive correlation between the expression of *MAD2* and *BUB1B* among the breast cancer-affected cases (Fig. 2F). Data from the representative cell lines of HR+ve background, as obtained from CCLE database, corroborated this observation (Supplementary file 2). Together, these data confirm that ERα transcriptionally regulates two of the MCC components, Mad2 and BubR1, in breast cancer cells.

### ER**α** influences the activity of mitotic checkpoint upon regulating MCC components in breast cancer cells

Our observation of ERα trans-regulating Mad2 and BubR1 led us to investigate the influence of the same on the progression of mitosis in breast cancer cells. Toward that, we synchronized HR+ve MCF-7 and HR-ve MDA-MB-231 cells with a double thymidine treatment at the G1/S transition. Following the release from second-time thymidine, subsets of cells were treated with the spindle poison, nocodazole, which induces mitotic arrest at the metaphase-anaphase transition (schematically shown in Fig. 3A). FACS analyses revealed mitotic exit of synchronized MCF-7 cells post 11 hours from the release of the G1-S block (Supplementary file 3, top panel and Supplementary table SI), while the addition of nocodazole resulted in mitotic blockage during that time-frame (Supplementary file 3, middle panel and Supplementary table I). In addition to this, the induction of cellular ERα activity, by 17β-estradiol (E2) treatment, also showed a similar observation (Supplementary file 3, bottom panel and Supplementary table I). Simultaneous WB analyses were performed to measure the level of a surrogate mitotic marker, Cyclin B1. The data confirmed nocodazole-imposed mitotic blockage in MCF-7 cells, as evidenced by the accumulation of Cyclin B1 (Fig. 3B, to compare the first three and next three lanes). A similar observation upon simultaneous E2 treatment confirms a functional mitotic arrest, irrespective of ligand addition in the media (Fig. 3B, last three lanes). On the contrary, despite the presence of nocodazole, cells of the MDA-MB-231 line showed a release from mitotic arrest, as apparent from FACS analyses (Supplementary file 4, top panel and Supplementary table II). Interestingly, the ectopic introduction of ERα brought about nocodazole-induced mitotic arrest in the MDA-MB-231 cells (Supplementary file 4, bottom panel and Supplementary table II). Further, WB data showed a decrease in Cyclin B1 in MDA-MB-231 cells even in the presence of nocodazole treatment (Fig. 3C, left panel). Interestingly, the level of Cyclin B1 was rescued when ERα was ectopically expressed in MDA-MB-231 cells (Fig. 3C, right panel). Concurrently, we knocked down the MCC component, Mad2, using a pool of siRNAs, to find out additional effects on the mitotic progression of these cells. Interestingly, we found that the siRNA-mediated downregulation of Mad2 resulted in a release of these cells from the mitotic arrest (Fig. 3D, right panel). These data, together, inform that ERα maintains the mitotic progression by transcriptionally activating the candidate MCC components and thus reports molecular crosstalk between ERα and mitosis in breast cancer cells. Furthermore, when we introduced ERα ectopically in MDA-MB-231 cells, we found a significant decrease in aneuploidy in these cells (Fig. 3E). This suggests an influence of ERα on the ploidy management of breast cancer cells.

**Figure 3:**
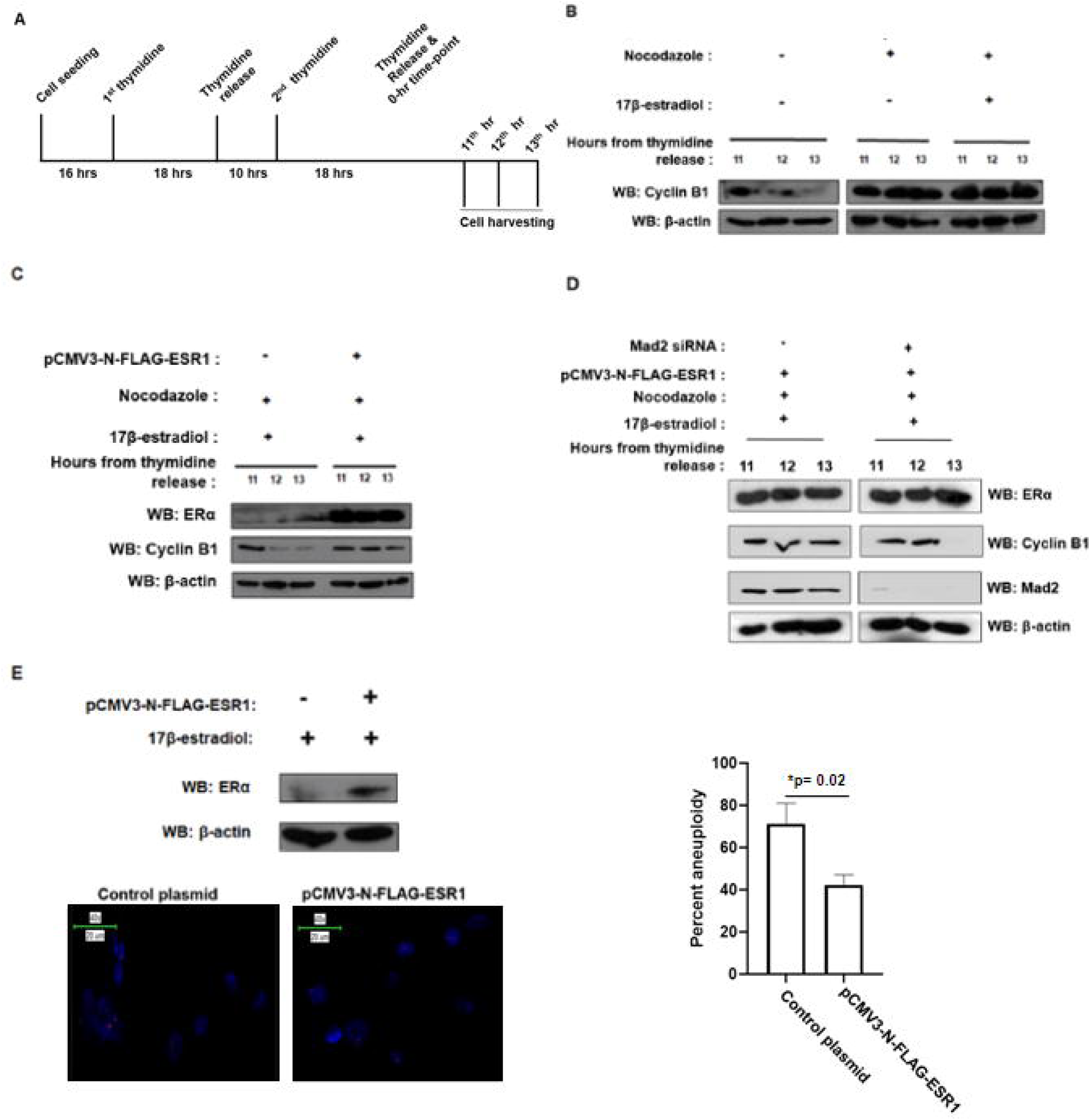
ERα regulates mitotic progression in breast cancer cells. A) Schematic representation of thymidine-mediated cell synchronization. B) Nocodazole arrests MCF-7 cells in mitosis. Cells were synchronized with double thymidine treatment (2.5 mM) as shown in (A) and subsequently, subsets of cells were treated with nocodazole (100 ng/ml) +/- 17-β-estradiol (10nM). Cells were harvested at indicated time points and processed for Western blot (WB) with indicated antibodies. C) Ectopic ERα restores mitotic blockage in MDA-MB-231 cells upon impairment of spindle assembly. Cells transiently transfected with either empty vector (pCMV-N-FLAG) or ERα-expressing pCMV-N-FLAG-ESR1 (1.5 µg) were synchronized as shown in (A) and treated with nocodazole (100 ng/ml) and 17-β-estradiol (10nM). Total cell lysates were processed for Western blot with indicated antibodies. D) siRNA-mediated downregulation of Mad2 resulted in a release of MDA-MB-231 cells from the mitotic arrest. Cells transiently transfected with ERα-expressing pCMV-N-FLAG-ESR1 (1.5 µg) were synchronized as shown in (A) and treated with nocodazole (100 ng/ml) and 17-β-estradiol (10nM). Additionally, these cells were transiently transfected with siRNA oligos (120nM) directed against Mad2 mRNA or scrambled siRNA oligo as control. Cells were harvested at indicated time points and processed for Western blot (WB) with indicated antibodies. E) The aneuploidy burden was reduced upon ectopic expression of ERα in MDA-MB-231 cells. Cells were processed for FISH and representative figures were shown. The experiment was performed in triplicate and the percent aneuploidies were plotted. The histograms show the percent aneuploidy in the Y axis. *p < 0.05; (two-tailed unpaired Student’s t-test).

### The level of MCC components influences survival outcomes in breast cancer

To investigate the clinical significance of *MAD2* and *BUB1B* in breast carcinoma-affected cases from the HR+ve background, we compared the expression of these two MCC components with clinicopathological variables upon retrieving data of the affected cases from the TCGA BRCA dataset. We segregated the samples into high and low *MAD2* or *BUB1B* expression groups based on the median of transcripts per million (TPM) values. To explore the influence of differential expression of these two genes on the clinicopathological features in the primary breast malignancies, we performed univariate analysis using the Chi-square test, as summarized in Supplementary Table III. Among the variables, the expression of the proliferation marker, Ki-67, showed a statistically significant association with both *MAD2* and *BUB1B* high groups (p <0.0001). Additionally, the *MAD2* high expression group showed a significant association with tumor stage (according to the American Joint Committee on Cancer guidelines), with a p-value of 0.0006. A similar observation was found in the case of high *BUB1B* expression groups and tumor stage (p < 0.0001). Although not reaching statistical significance, a close association was observed in the case of tumor nodal involvement for the high *MAD2* expression group (p = 0.0574). However, no statistically significant associations were found between the expression groups of *BUB1B* and nodal involvement. Additionally, different histologic subtypes of breast tumors, as per ICD-0-3 guidelines, namely intraductal papillary carcinoma with invasion, pleomorphic carcinoma, adenoid cystic carcinoma, cribriform carcinoma, and infiltrating lobular mixed carcinoma showed significant greater occurrence in high-expressing groups of these two MCC genes (p < 0.0001).

To investigate the effect of different clinicopathological variables on recurrence free survival and overall survival, we included them in a multivariate Cox proportional hazards regression model (Supplementary Table IV). In this multivariable analysis, we found that high Ki67 expression was associated with both poor OS (HR= 4.566, p<0.001) and RFS (HR= 4.514, p<0.001). Similarly, the histological type of IDC was also associated with both poor OS (HR=1.281, p=0.025) and RFS (HR=1.225, p=0.035). In the case of high *BUB1B* expression, the hazard ratio was similar to the reference category of low *BUB1B* expression and the p-value was not significant. In the case of high *MAD2* expression, we observed that the hazard ratio was 1.242 (OS) and 1.226 (RFS), respectively, although the p values were non-significant.

In the prospective study cohort of Eastern Indian origin, we analyzed the clinicopathological features of the individuals diagnosed with HR+ve primary breast malignancies. The cases were categorized into groups based on the median value of gene expression of *MAD2* or *BUB1B* and named ‘*MAD2* High’, ‘*MAD2* Low’ ‘*BUB1B* High’, and ‘*BUB1B* Low’. We performed univariate analysis using the Chi-square test, as summarized in Supplementary Table V, to explore the association of these two genes with different clinicopathological features (i.e., stage, grade, nodal involvement, histologic type, Ki-67 level, lymphovascular invasion/LVI and perineural invasion/ PNI). Among the variables examined, the expression of the proliferation marker Ki-67 exhibited a significant association with the high expression group of both *MAD2* (p=0.0035) and *BUB1B* (p=0.0477). The tumor stage demonstrated significant associations with the high *MAD2* (p < 0.0001) as well as *BUB1B* expression groups (p < 0.0001). Similarly, tumor grade showed significant associations with the high *MAD2* as well as *BUB1B* expression groups (p < 0.0001).

Considering the nodal involvement, statistical significance was found in the case of the high *MAD2* group (p < 0.0001). However, no statistically significant associations were found between the expression groups of *BUB1B* and nodal involvement. Histologic type exhibited significantly greater occurrence in high-expressing groups of both of the genes (p < 0.0001). LVI and PNI are recognized as indicators of poor outcomes in various cancers, including breast cancer. LVI did not show a significant association with *MAD2* expression but demonstrated a significant association in the high *BUB1B* expression group (p=0.0004). No statistical significance was observed for PNI with *MAD2* or *BUB1B* expression.

We further elaborated our findings of differential expression of *MAD2* or *BUB1B* with the survival outcome among breast cancer sufferers. We segregated the cases of HR+ve breast cancer from the TCGA cohort into two groups, based on the transcript levels of *MAD2* or *BUB1B*. The ‘*MAD2* high’ expressing group showed poorer overall survival, compared to the ‘*MAD2* low’ expressing group (Fig. 4A, left panel). A similar observation was found when the cases were analyzed for that of *BUB1B* expression (Fig. 4A, right panel). For the analysis of recurrence free survival in the same cohort, *MAD2* (Fig. 4B, left panel) as well as *BUB1B* high-expressing groups (Fig. 4B, right panel) exhibited significantly poorer outcome. Similarly, we segregated our prospective study cohort of Eastern Indian cases. Though not significant, a similar trend was observed in their survival outcome when analyzed based on *MAD2* (Supplementary Figures 5 and 6) or *BUB1B* expression (Supplementary Figures 5 and 6). As supporting evidence, we performed a clonogenic assay under the influence of ectopic Mad2. We observed that cells from the HR+ve breast cancer line, T-47D, showed higher colony numbers with an upregulation of Mad2 expression (Fig. 4C), indicating the effect of deregulated MAD2 in influencing the overall proliferative capabilities of the breast cancer cell line. Together, these data support that ERα-driven trans-regulation of MCC components, Mad2 and BubR1, contributes to the poor survival outcome among the affected cases of breast cancer.

**Figure 4:**
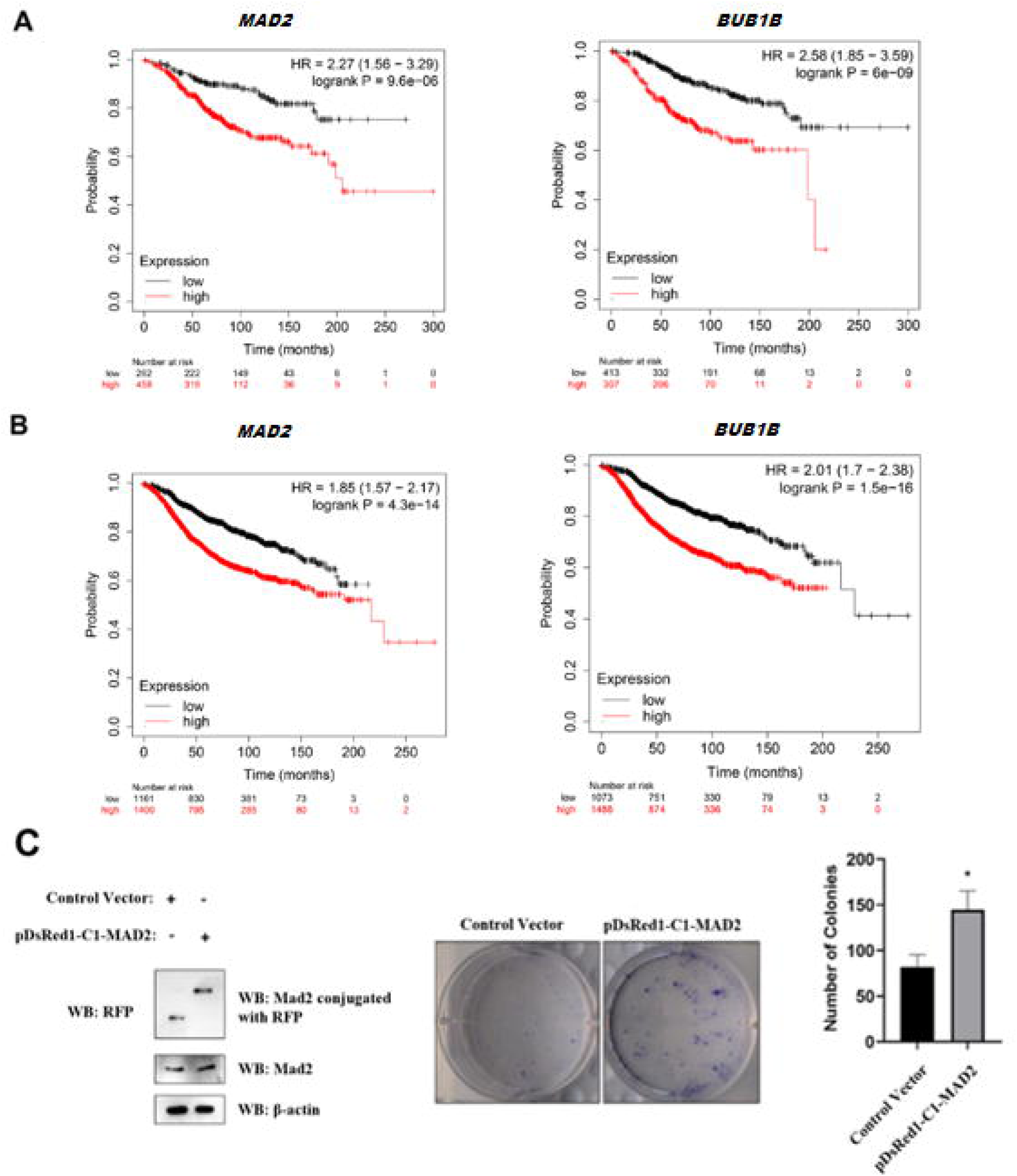
Prognostic significance of *MAD2* and *BUB1B* in primary breast malignancies. A) Kaplan-Meier analysis to detect the overall survival of HR+ve breast cancer-affected cases in TCGA BRCA cohort, according to their expression of *MAD2* and *BUB1B* (high vs low). Log rank P *< 0.05 B) Kaplan-Meier analysis to detect the recurrence free survival of HR+ve breast cancer-affected cases in TCGA BRCA cohort, according to their expression of *MAD2* and *BUB1B* (high vs low). Log rank P *< 0.05 C) Upregulated Mad2 expression promotes clonogenicity in breast cancer cells. T-47D cells were transiently transfected with either an empty vector or Mad2-expressing plasmid, pDsRed1-C1-MAD2 (1µg). After 48 hours of transfection, a part of the cells was seeded in 6-well plates and incubated for one week to obtain visible colonies. Simultaneously, the transfection is validated by Western blot from another part of the cells by using antibodies specific for RFP, Mad2 and, β-actin. Colonies were stained with crystal violet and subsequently counted. Data was expressed as mean ± SD, represented as a histogram. Colonies were stained with crystal violet and subsequently counted. Data was expressed as mean ± SD, represented as a histogram. All the experiments were performed thrice and the Student’s t-test was used to analyze the data. *p < 0.05.

## Discussion

ERα plays a crucial role in the growth and maintenance of the mammary gland and is responsible for promoting around 70% of breast cancer cases [1]. ERα is a transcription factor, which upon activation by its ligand, estrogen, binds to chromatin, modulating gene expression. Importantly, ERα is involved in interactions with Cyclin A and Cyclin D1, which promotes cell division by regulating phosphorylation of the retinoblastoma protein (pRb) [52, 53]. Other ER-responsive genes include c-MYC and CDK inhibitor p27, both of which are also involved in cell cycle progression [54, 55]. Reportedly, a G2/M kinase named PLK1 (polo-like kinase 1) interacts with ERα and is recruited to EREs, thus coactivating the transcription of ERα-target genes in breast cancer cells [56]. A prior study also documented a direct interaction of another hormone receptor member, estrogen receptor β (ERβ) with the MCC component, Mad2, thus promoting cellular proliferation and regulation of the cell cycle [57]. Forwarding these observations, the results of our study could highlight a direct trans-regulatory role of ERα on the two MCC proteins, BubR1 and Mad2, both of which are involved in the regulation of chromosome segregation during mitosis. To note, defects in the mitotic checkpoint, reportedly, lead to aneuploidy, promoting malignancy [58, 59]. Such fate of cells is often attained upon the inability to execute the programmed cell death mechanism, apoptosis. Reports showed that losing the ability to commit apoptosis under such conditions contributes to the development of cancer [60]. Notably, cells with an impaired mitotic checkpoint experience less apoptotic cell death compared to cells with a functional checkpoint, emphasizing the necessity of a well-functioning mitotic checkpoint for apoptosis induction [61]. Recent studies, more specifically, investigated the role of BubR1 and Mad2 in aneuploidy and breast cancer. Haploinsufficiency of BubR1 promotes the formation of micronuclei and aneuploidy, resulting in tumor formation in mice [62, 63]. Another report demonstrated that transient Mad2 overexpression promotes chromosomal instability and aneuploidy, thereby stimulating the initiation and progression of cancer in mice [64]. Towards this, our study could add a new lead on the aberrant levels of BubR1 and Mad2 behind the poor outcome of breast cancer. To note, a prior report showed that *MAD2* exhibits putative estrogen and progesterone response elements, on its promoter [48]. Our study attempted to frame a comprehensive picture of the role of ERα in regulating mitosis in breast cancer. We focused on the MCC machinery and the initial clues came from the TCGA BRCA cohort as well as a prospective breast cancer cohort of Eastern Indian cases. Both cohorts revealed higher levels of two of the MCC components, Mad2 and BubR1 in tumor specimens compared to the adjacent normal tissues. Further segregation of the prospective study cohort showed a higher expression of Mad2 and BubR1 among HR+ve cases. Cultured cells from HR+ve lines also replicated this observation. This made us curious and we selected ERα as the representative member of HR for the next sets of experiments. Results from these experiments could show a positive influence of ERα on the expression of Mad2 and BubR1. Considering the transcriptional activity of ERα, we went further and decoded a direct role of ERα in the trans-regulation of these two genes. We also correlated this ERα-driven transcriptional program in imparting a poor survival outcome in both the TCGA BRCA and the prospective study cohort and further could link this observation with the higher clonogenic potential of Mad2 in our experimental set-up. We would mention that, in some instances of the prospective cohort, when we compared the transcript levels or survival outcomes, we could not reach the level of statistical significance. However, we could not rule out the influence of the limited cohort size or assessment period (less than five years), for the same. Nonetheless, this study could find two direct targets of ERα, both of which are major contributors to mitotic progression and ploidy regulation during cell division. To note, the third MCC component, Bub3, did not show similar effects, in this respect. However, we anticipate an indirect effect of HR on Bub3, as HR+ve cells generally showed higher Bub3 levels in our experimental set-up.

Previous study from our lab revealed that the hormone-receptor-degrading-protein, CUEDC2 modulates cellular Mad2 and BubR1 levels in breast cancer, in an ERα-dependent manner [49]. Moreover, this CUEDC2-ERα crosstalk showed an influence on the ploidy outcome of breast cancer cells by regulating mitosis. The present study furthered this by finding the direct role of ERα in trans-regulating Mad2 and BubR1. Towards this, deregulated levels of these proteins reportedly contribute to aneuploidy in various cancers [62–65]. Aneuploidy is one of the prominent features among CIN phenotypes and is observed in more than 80% of cases of malignant solid tumors [66]. The identification of methods that utilize chromosomal instability (CIN) to specifically target cancer cells is a complex area of study in cancer biology, holding significant implications for the advancement of anti-cancer drug therapies. In transgenic mice with SAC deficiency, upregulation of core SAC components like Bub1 or Mad2, exhibit alterations in the number of copies of entire chromosomes, as well as an increased propensity for tumor development. [64, 67]. As SAC disruption and chromosomal instability (CIN) are deeply linked, MCC components are promising targets for therapeutic advancement. Currently, two primary therapeutic approaches exist for leveraging CIN in cancer treatment, commonly known as CIN-reducing and CIN-inducing strategies. CIN-reducing treatments aim to inhibit the formation of further CIN in tumors with existing chromosomal instability, to decrease or prevent further genetic abnormalities. Ideally, this would impede the ability of tumors to adapt, restrict the evolution of cancer cells, and prevent the development of drug resistance, resulting in a decrease in tumor aggressiveness. On the contrary, CIN-inducing strategies aim to induce higher numerical and/or structural chromosomal aberrations to surpass a certain threshold, triggering programmed cell death [68]. Together, investigating and harnessing CIN in cancer holds significant therapeutic promise and could serve as a crucial vulnerability to be targeted for the treatment of aggressive and drug-resistant cancers, ultimately leading to improved quality of life and better outcome for individuals affected by cancer. In this respect, we propose this and other transcriptional programs of ERα on cell cycle regulation as potential alleys to dissect further.

## Conclusions

Our study revealed that ERα transcriptionally upregulates two of the MCC components, Mad2 and BubR1, thus influencing the mitotic progression of breast cancer cells. This ERα-driven transcriptional upregulation showed an association with the survival outcome of breast cancer sufferers from the TCGA BRCA and an Eastern Indian study cohort. Further elucidation of this molecular pathway holds translational potential for the clinical management of breast cancer.

## List of abbreviations

ERα: Estrogen Receptor alpha
MCC: Mitotic Checkpoint Complex
HR: Hormone Receptor
PR: Progesterone Receptor
HR+ve: HR positive
HER2: Human Epidermal growth factor Receptor 2
TNBC: Triple Negative Breast Cancer
SAC: Spindle Assembly Checkpoint
APC/C: Anaphase-Promoting Complex/Cyclosome
CIN: Chromosomal Instability
TCGA BRCA: The Cancer Genome Atlas breast invasive carcinoma
ERE: Estrogen Response Element
CCLE: Cancer Cell Line Encyclopedia
ERβ: Estrogen Receptor β
Bub3: Budding Uninhibited by Benzimidazoles 3
Mad2: Mitotic arrest deficient 2
BubR1: Budding uninhibited by benzimidazole-Related 1

## Declarations

### Ethical approval and consent to participate

Primary malignant breast tumor and adjacent normal tissues were collected during surgery with written informed consent from all the participants or their kins at Saroj Gupta Cancer Centre and Research Institute (SGCC&RI), Kolkata. The study was approved by the institutional ethics committee of Saroj Gupta Cancer Centre and Research Institute (IEC SGCCRI REF NO-2016/3/4/SN/NON-REG/05/02) under the regulation of the Indian Council of Medical Research (ICMR), Govt. of India (Registration No. ECR/250/Inst/WB/2013/RR-20). All methods were carried out according to relevant guidelines and regulations.

### Consent for publication

Not applicable

### Availability of data and materials

The gene expression data were extracted from the TCGA BRCA dataset from the UALCAN database (https://ualcan.path.uab.edu/). The dataset containing the gene expression data analyzed during the current study is available in the 22Q2 public data repository from the Depmap portal at Broad Institute, https://depmap.org/portal/). The Kaplan-Meier plotter database (http://kmplot.com/analysis/) site was used to analyze the survival outcome employing ‘203362_s_at’ and ‘203755_at’ datasets.

### Authors’ contributions

CONCEPTION: SS, SR, and, SN conceptualized the manuscript.

DATA CURATION: SS, RP, DD and AM helped in data curation. SS, RP, DD and, SR performed the experiments.

ANALYSIS OF DATA: SS, RP, AM, DD, and, SN analyzed the data.

PREPARATION OF THE MANUSCRIPT: SS, DD, SR, RP, and SN wrote the manuscript with input from all the authors.

REVISION FOR IMPORTANT INTELLECTUAL CONTENT: All authors contributed to the article and approved the submitted version.

SUPERVISION: SN supervised all the experiments performed and also the analysis of the data.

## Acknowledgments

We acknowledge Prof. Tanya Das, Bose Institute, Kolkata, India, and, Dr. Snehasikta Swarnakar, Indian Institute of Chemical Biology, Kolkata, India, for their cell culture-related help. We acknowledge Dr. Chandrama Mukherjee, Institute of Health Sciences, Presidency University, Kolkata, India for kindly providing the wild-type construct of RFP. We acknowledge Rajeswari Dutta, SGCCRI, Kolkata, India, for FACS-related support. We also acknowledge Prof. Susanta Roychoudhury, ICMR Emeritus Scientist, CSIR-Indian Institute of Chemical Biology, Kolkata, India for his valuable input.

## Authors’ information

SS, SR, RP, AM, SK, AG, and, DD are associated with the Department of Basic and Translational Research, SGCCRI, Kolkata, India. DD and SN are currently affiliated with the Institute of Health Sciences, Presidency University, Kolkata, India.

