## Supplementary File for "Estrogen receptor alpha (ERα) driven trans-regulation of mitotic checkpoint complex (MCC) components affects the clinical outcome of breast cancer"

**Supplementary files**

The estrogen treatment resulted in a significant increase of luciferase activity from the pSS-BUB1B as well as pSS-MAD2 constructs when transfected in HR+ve breast cancer line, T-47D (Supplementary figure 1). Data from the representative cell lines of HR+ve background, as obtained from the CCLE database, showed a positive correlation between the expression of *MAD2* and *BUB1B* (Supplementary figure 2). Flow cytometry analysis from the synchronized MCF-7 cells revealed a functional G2/M arrest imposed by nocodazole (Supplementary Figure 3 and Supplementary Table I). Additionally, in the case of MDA-MB-231 cells, flow cytometry depicted that Ectopic ERα restores G2/M arrest upon impairment of spindle assembly (Supplementary Figure 4 and Supplementary Table II). The association between *MAD2* and *BUB1B* expression with clinicopathological features in breast cancer-affected individuals from the TCGA BRCA cohort was shown in Supplementary Table III. Further, Multivariate Cox regression analysis of overall survival and recurrence free survival was depicted in Supplementary Table IV on different clinicopathological variables in the TCGA BRCA cohort. The association between *MAD2* and *BUB1B* expression with clinicopathological features in breast cancer-affected individuals from the Eastern Indian cohort was shown in Supplementary Table V. Moreover, Kaplan-Meier analysis was performed to detect the overall as well as recurrence free survival of HR+ve breast cancer-affected cases in the prospective study cohort of Eastern Indian individuals, according to their expression of *MAD2* and *BUB1B* (high vs low) (Supplementary Figures 5 and 6).

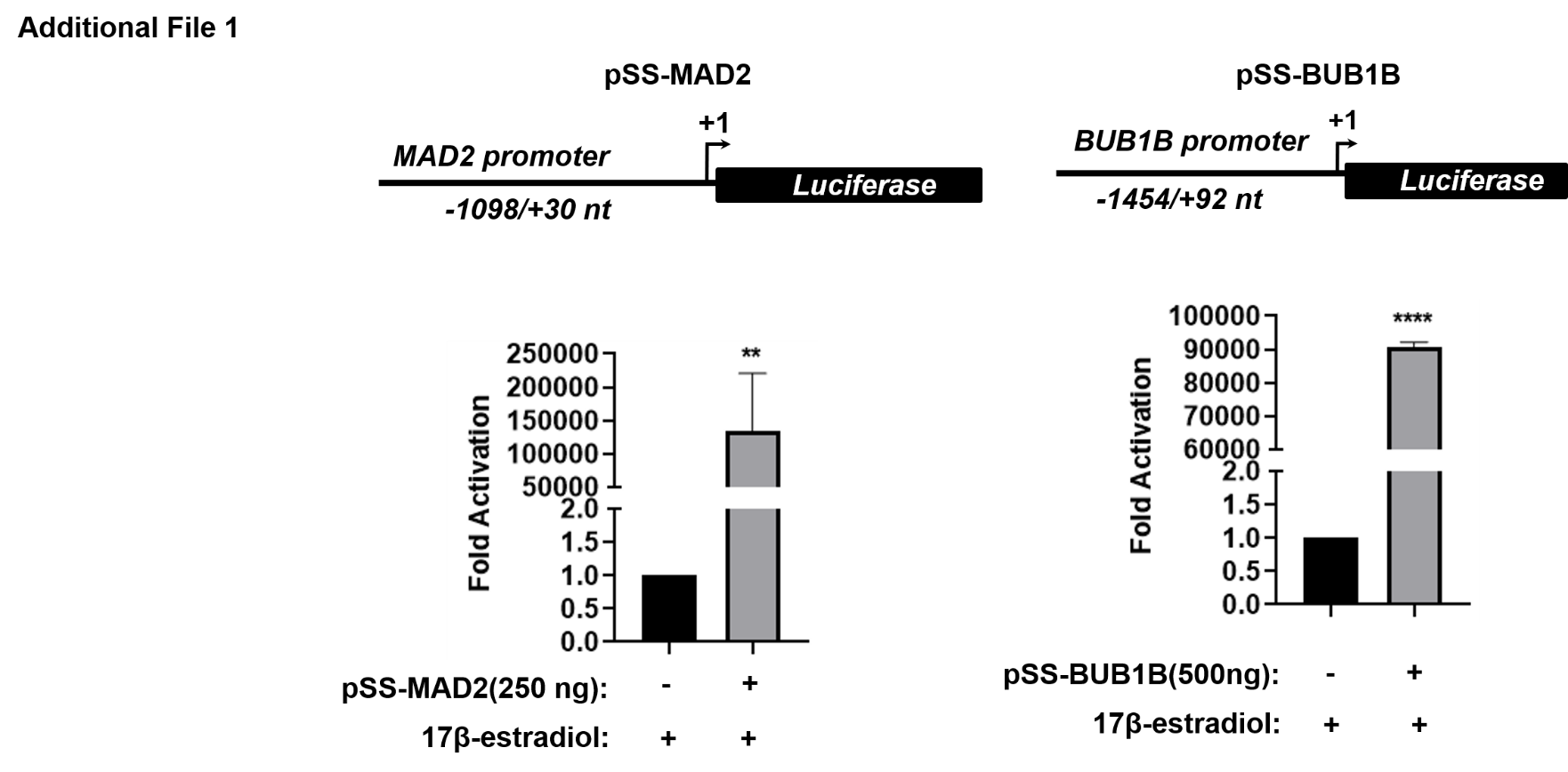

**Supplementary Figure 1:** E2 (17β-estradiol) treatment up-regulates *MAD2* and *BUB1B* promoter activity in T-47D cells. Top panel shows the map of *MAD2* and *BUB1B* promoter regions used for luciferase reporter assay. pSS-MAD2 construct contains a 1128-bp region (-1098 to +30 nt) of *MAD2* gene and pSS-BUB1B construct contains a 1546-bp region (-1454 to +92 nt) of *BUB1B* gene, cloned into pGL3 basic vector. T-47D cells were transiently transfected with pSS-MAD2 (250 ng) and pSS-BUB1B (500 ng), followed by E2 treatment and proceeded for luciferase reporter assay. **p < 0.005; ***p < 0.0005; (two-tailed unpaired Student’s t-test)

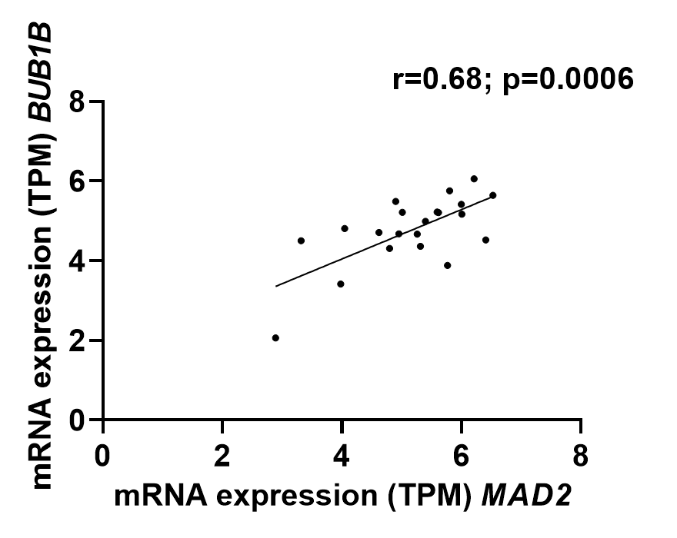

**Supplementary Figure 2:** Representative scatter plot depicting the Pearson’s correlation analysis between the gene expression values (TPM) of *MAD2* and *BUB1B*, extracted from CCLE database. The Pearson correlation coefficient is reported as ‘r’ value. *******p < 0.0005; (two-tailed unpaired Student’s t-test).

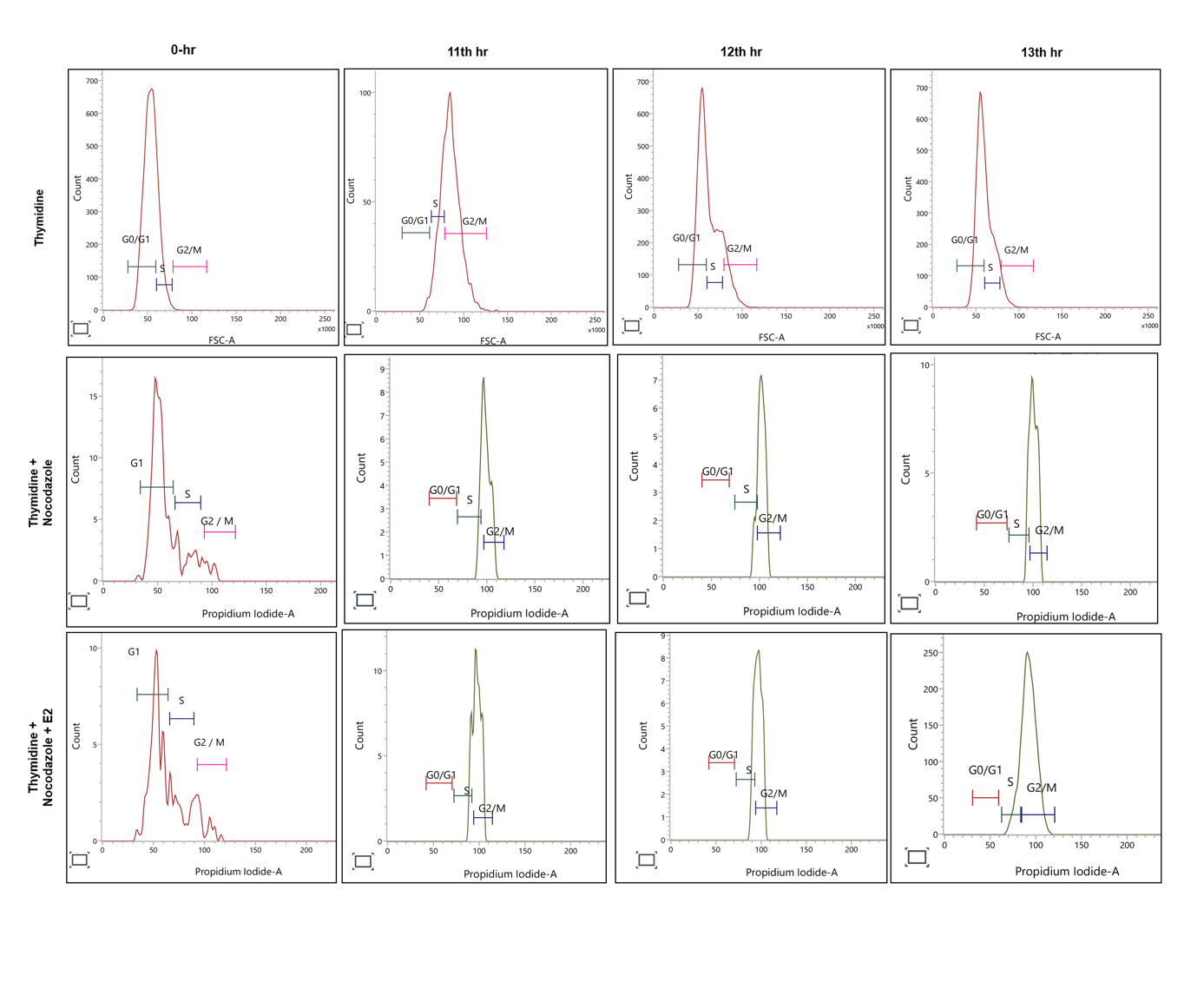

**Supplementary Figure 3:** Nocodazole imposes G2/M arrest in MCF-7 cells. Cells were synchronized with double thymidine treatment (2.5 mM) and subsequently, subsets of cells were treated with nocodazole (100 ng/ml) +/- 17-β-estradiol (10nM). Cells were harvested at indicated time points and processed for flow cytometry.

| Hours from thymidine release  Percentage  of cells | 0-hr | | | 11^th^ hr | | | 12^th^ hr | | | 13^th^ hr | | |
| --- | --- | --- | --- | --- | --- | --- | --- | --- | --- | --- | --- | --- |
|  | G0/G1  (%) | S  (%) | G2/M  (%) | G0/G1  (%) | S  (%) | G2/M  (%) | G0/G1  (%) | S  (%) | G2/M  (%) | G0/G1  (%) | S  (%) | G2/M  (%) |
| Thymidine | 74.91 | 24.86 | 0.23 | 1.17 | 24.29 | 74.54 | 51.2 | 34.14 | 14.67 | 53.8 | 40.59 | 5.61 |
| Thymidine + Nocodazole | 77.85 | 16.6 | 5.54 | 0 | 13.33 | 86.67 | 0 | 11.84 | 88.15 | 0 | 13.73 | 86 |
| Thymidine+Nocodazole+E2 | 65.43 | 23.94 | 10.64 | 0 | 14.51 | 85.48 | 0 | 22.77 | 77.23 | 0 | 12.82 | 87.18 |

**Supplementary Table I:** The percentage of MCF-7 cells at different cell cycle phases

| Hours from thymidine release  Percentage  of cells | 0-hr | | | 11^th^ hr | | | 12^th^ hr | | | 13^th^ hr | | |
| --- | --- | --- | --- | --- | --- | --- | --- | --- | --- | --- | --- | --- |
|  | G0/G1  (%) | S  (%) | G2/M  (%) | G0/G1  (%) | S  (%) | G2/M  (%) | G0/G1  (%) | S  (%) | G2/M  (%) | G0/G1  (%) | S  (%) | G2/M  (%) |
| Nocodazole + Empty vector + E2 | 48.65 | 51.35 | 0 | 0 | 0 | 99.84 | 58.69 | 16.2 | 25.12 | 65.22 | 32.33 | 2.46 |
| Nocodazole + Ectopic ERα + E2 | 94.11 | 5.88 | 0 | 0 | 0 | 100 | 10.64 | 24.82 | 64.54 | 39.73 | 35.96 | 24.32 |

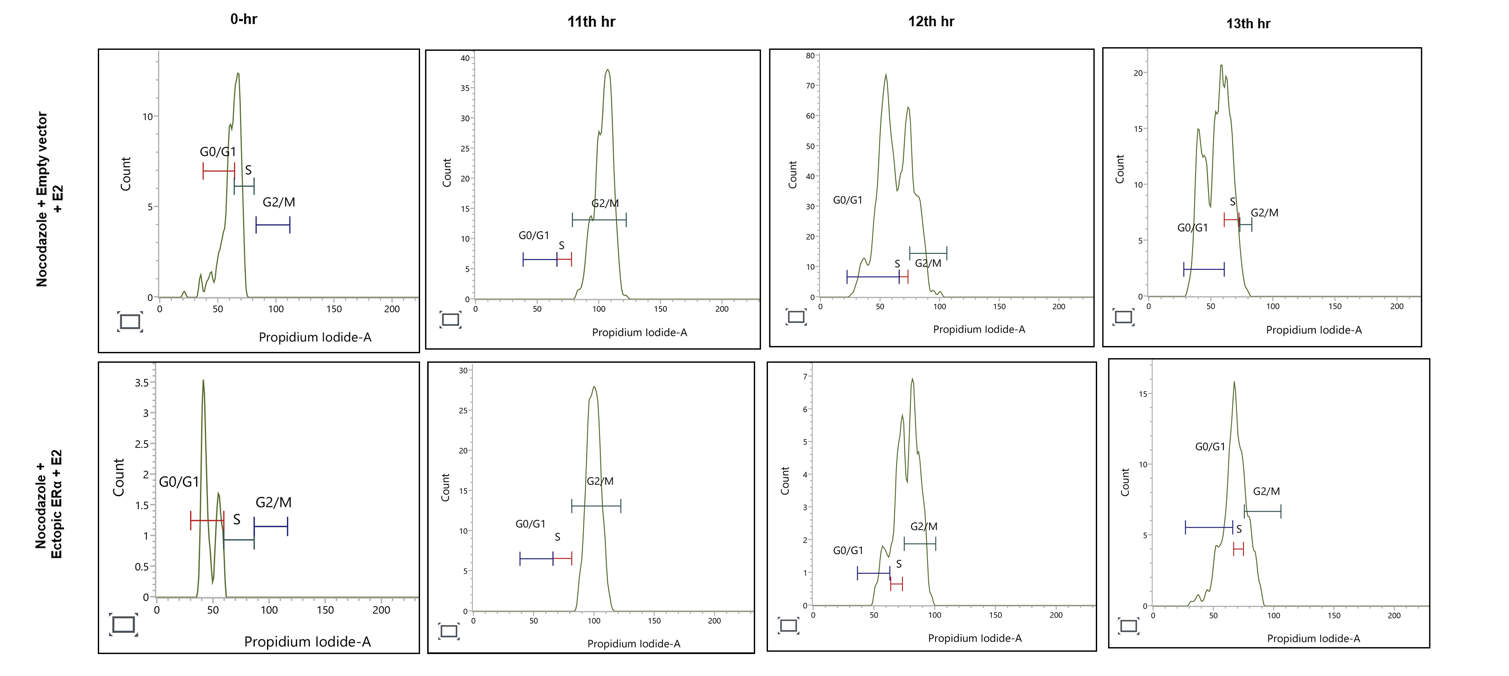

**Supplementary Figure 4:** Ectopic ERα restores G2/M arrest in MDA-MB-231 cells upon impairment of spindle assembly. Cells, transfected with either empty vector (pCMV-N-FLAG) or ERα-expressing pCMV-N-FLAG-ESR1 (1.5 µg) were synchronized with double thymidine treatment (2.5 mM) and treated with nocodazole (100 ng/ml) and 17-β-estradiol (10nM) prior to harvesting for flow cytometry.

**Supplementary Table II:** The percentage of MDA-MB-231 cells at different cell cycle phases

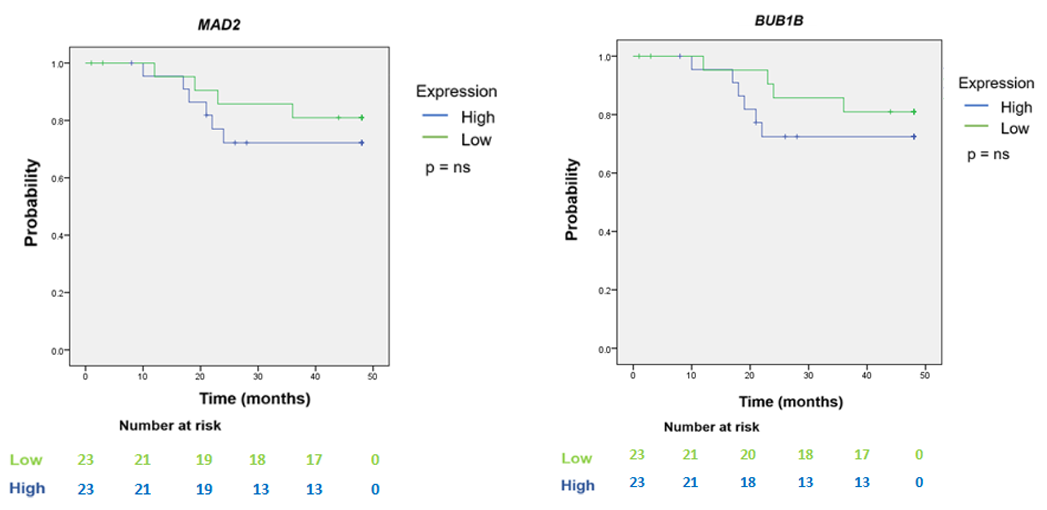

**Supplementary Figure 5:** Kaplan-Meier analysis to detect the overall survival of HR+ve breast cancer-affected cases in the prospective study cohort of Eastern Indian individuals, according to their expression of *MAD2* and *BUB1B* (high vs low). ns, not significant.

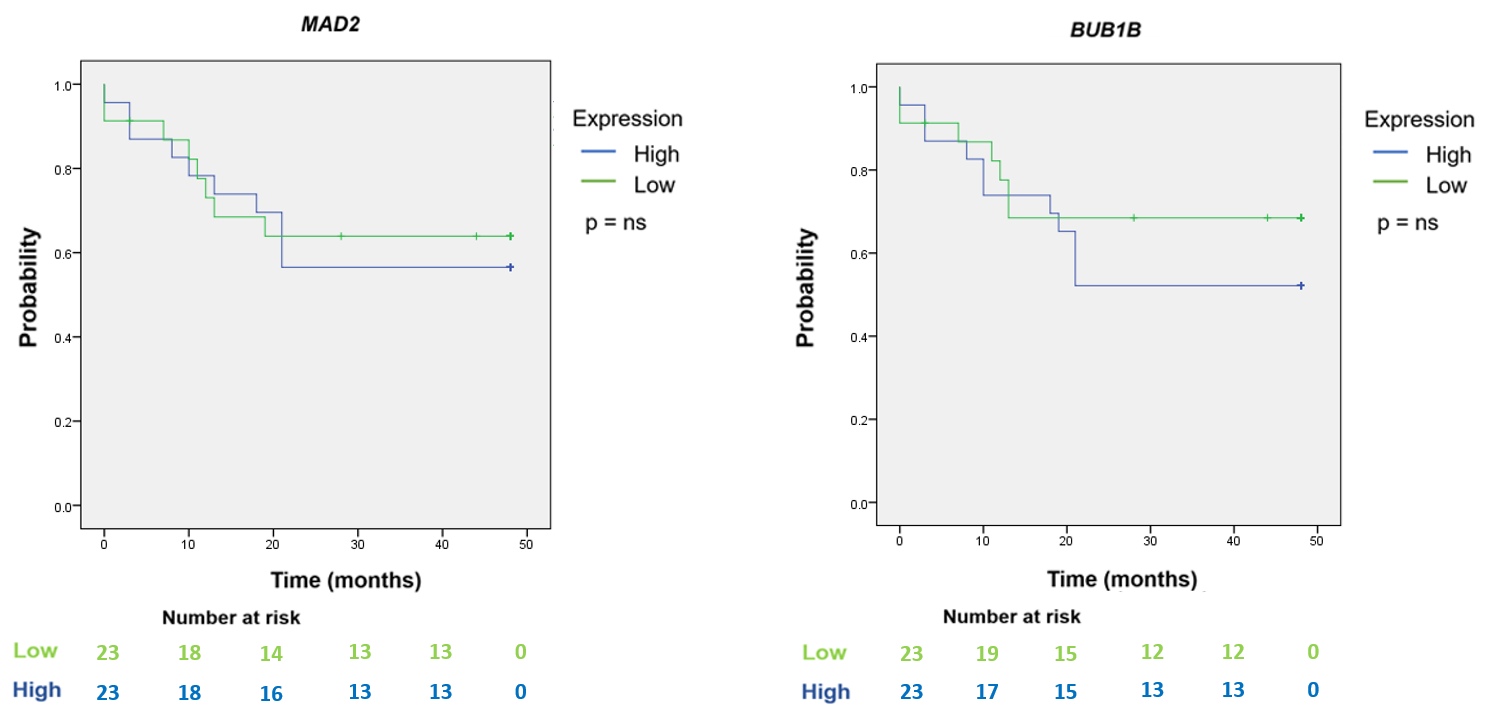

**Supplementary Figure 6:** Kaplan-Meier analysis to detect the recurrence free survival of HR+ve breast cancer affected cases in the prospective study cohort of Eastern Indian individuals, according to their expression of *MAD2* and *BUB1B* (high vs low). ns, not significant.

| ***Variables (TCGA BRCA Cohort)*** | ***MAD2 expression*** | | | ***BUB1B expression*** | | |
| --- | --- | --- | --- | --- | --- | --- |
|  | ***Low N (%)*** | ***High N (%)*** | ***p-Value^a^*** | ***Low N (%)*** | ***High N (%)*** | ***p-Value^a^*** |
| Stage  I  II  III  IV | 39 (37.1)  159 (48.7)  84 (61.7)  8 (61.5) | 66 (62.8)  167 (51.2)  52 (38.2)  5 (38.4) | 0.0006 | 41 (39)  164 (57.3)  77 (56.6)  8 (80) | 64 (61)  122 (42.6)  59 (43.3)  2 (20) | <0.0001 |
| Nodal Involvement  N_0_  N_1_  N_2_  N_3_ | 120 (44.4)  105 (51.7)  48 (61.5)  17 (58.6) | 150 (55.5)  98 (48.2)  30 (38.4)  12 (41.3) | 0.0574 | 125 (46.2)  100 (49.2)  49 (62)  16 (55.1) | 145 (53.7)  103 (50.7)  30 (38)  13 (44.8) | 0.1106 |
| Histological Type*  IDC  ILC  DLC  MC  IDP with invasion  IDC Mixed  PD  MC  PC  ACC  CC  ILMC | 245 (51.2)  17 (26.9)  12 (52.1)  3 (60)  1 (33.3)  8 (53.3)  2 (100)  1 (100)  0 (0)  0 (0)  0 (0)  0 (0) | 214 (48.7)  46 (73)  11 (47.8)  2 (40)  2 (66.6)  7 (46.6)  0 (0)  0 (0)  1 (100)  1 (100)  1 (100)  2 (100) | <0.0001 | 244 (53.1)  22 (36)  11 (47.8)  2 (40)  1 (33.3)  6 (40)  2 (100)  1 (100)  0 (0)  0 (0)  0 (0)  0 (0) | 215 (46.8)  41 (64)  12 (52.1)  3 (60)  2 (66.6)  9 (60)  0 (0)  0 (0)  1 (100)  1 (100)  1 (100)  1 (100) | <0.0001 |
| Ki67  High  Low | 368 (80.7)  88 (19.3) | 88 (19.3)  367 (80.6) | <0.0001 | 388 (85)  68 (14.9) | 68 (15)  387 (85) | <0.0001 |

*IDC: Infiltrating Ductal Carcinoma, ILC: Infiltrating Lobular Carcinoma, DLC: Ductal Lobular Carcinoma, MC: Mucinous Adenocarcinoma, IDP: Intraductal Papillary, PD: Paget Disease, MC: Metaplastic Carcinoma, PC: Pleomorphic Carcinoma, ACC: Adenoid Cystic Carcinoma, CC: Cribriform carcinoma, ILMC: Infiltrating Lobular Mixed Carcinoma

**^a^**Chi-Square Test

**Supplementary Table III:** Association between *MAD2* and *BUB1B* expression with clinicopathological features in breast cancer-affected individuals from TCGA BRCA cohort

| ***Overall Survival*** | | | ***Recurrence Free Survival*** | |
| --- | --- | --- | --- | --- |
| ***Variables*** | ***Hazard Ratio (95% CI)*** | ***p-Value*** | ***Hazard Ratio (95% CI)*** | ***p-Value*** |
| **Stage**  Low (I, II)  High (III, IV) | 1.00  1.044 (0.771-1.415) | 0.780 | 1.00  0.989 (0.732-1.336) | 0.942 |
| **Nodal Involvement**  Low (N0, N1)  High (N2, N3) | 1.00  1.152 (0.819-1.621) | 0.416 | 1.00  1.183 (0.844-1.657) | 0.329 |
| **Ki67 Expression**  Low  High | 1.00  4.566 (3.724-5.598) | <0.001 | 1.00  4.514 (3.688-5.526) | <0.001 |
| **Histological Type**  Others  Infiltrating Ductal Carcinoma (IDC) | 1.00  1.281 (1.032-1.589) | 0.025 | 1.00  1.255 (1.016-1.551) | 0.035 |
| ***BUB1B* Expression**  Low  High | 1.00  1.065 (0.845-1.343) | 0.592 | 1.00  1.007 (0.805-1.261) | 0.949 |
| ***MAD2* expression**  Low  High | 1.00  1.242 (0.929-1.660) | 0.143 | 1.00  1.226 (0.837-1.318) | 0.673 |

**Supplementary Table IV:** Multivariate Cox regression analysis of Overall survival and Recurrence free survival on different clinicopathological variables in TCGA BRCA cohort

| ***Variables (Eastern Indian Cohort)*** | ***MAD2 expression*** | | | ***BUB1B expression*** | | |
| --- | --- | --- | --- | --- | --- | --- |
|  | ***Low N (%)*** | ***High N (%)*** | ***p-Value^a^*** | ***Low N (%)*** | ***High N (%)*** | ***p-Value^a^*** |
| Stage  I  II  III  IV | 1 (100)  13 (50)  5 (41.6)  1 (33.3) | 0 (0)  13 (50)  7 (58.3)  2 (66.6) | <0.0001 | 1 (100)  12 (46.1)  5 (41.6)  1 (33.3) | 0 (0)  14 (53.8)  7 (58.3)  2 (66.6) | <0.0001 |
| Grade  G1  G2  G3 | 0 (0)  7 (43.7)  12 (54.5) | 1 (100)  9 (56.2)  10 (45.4) | <0.0001 | 0 (0)  9 (56.2)  9 (42.8) | 1 (100)  7 (43.7)  12 (57.1) | <0.0001 |
| Nodal Involvement  N_0_  N_1_  N_2_  N_3_ | 7 (53.8)  7 (50)  2 (28.5)  3 (60) | 6 (46.1)  7 (50)  5 (71.4)  2 (40) | <0.0001 | 6 (46.1)  7 (50)  3 (42.8)  2 (40) | 7 (53.8)  7 (50)  4 (57.1)  3 (60) | 0.5291 |
| Histological Type*  IDC  ILC  IDC-Mucinous | 21 (51.2)  0 (0)  0 (0) | 20 (48.7)  1 (100)  2 (100) | <0.0001 | 21 (51.2)  0 (0)  0 (0) | 20 (48.7)  1 (100)  2 (100) | <0.0001 |
| Ki67  High  Low | 17 (48.5)  2 (28.5) | 18 (51.4)  5 (71.4) | 0.0035 | 4 (57.1)  15 (42.8) | 3 (42.8)  20 (57.1) | 0.0477 |
| Lymphovascular invasion (LVI)  Present  Absent | 10 (50)  8 (47) | 10 (50)  9 (53) | 0.6712 | 12 (60)  6 (35.2) | 8 (40)  11 (64.7) | 0.0004 |
| Perineural invasion  Present  Absent | 4 (50)  14 (48.2) | 4 (50)  15 (51.7) | 0.7773 | 4 (50)  14 (48.2) | 4 (50)  15 (51.7) | 0.7773 |

*IDC: Infiltrating Ductal Carcinoma, ILC: Infiltrating Lobular Carcinoma, DLC: Ductal Lobular Carcinoma, MC: Mucinous Adenocarcinoma, IDP: Intraductal Papillary, PD: Paget Disease, MC: Metaplastic Carcinoma, PC: Pleomorphic Carcinoma, ACC: Adenoid Cystic Carcinoma, CC: Cribriform carcinoma, ILMC: Infiltrating Lobular Mixed Carcinoma

**^a^**Chi-Square Test

**Supplementary Table V:** Association between *MAD2* and *BUB1B* expression with clinicopathological features in breast cancer-affected individuals from the Eastern Indian cohort
